# Automation of Leaf Counting in Maize and Sorghum Using Deep Learning

**DOI:** 10.1101/2020.12.19.423626

**Authors:** Chenyong Miao, Alice Guo, Addie M. Thompson, Jinliang Yang, Yufeng Ge, James C. Schnable

## Abstract

Leaf number and leaf emergence rate are phenotypes of interest to plant breeders, plant geneticists, and crop modelers. Counting the extant leaves of an individual plant is straightforward even for an untrained individual, but manually tracking changes in leaf numbers for hundreds of individuals across multiple time points is logistically challenging. This study generated a dataset including over 150,000 maize and sorghum images for leaf counting projects. A subset of 17,783 images also includes annotations of the positions of individual leaf tips. With these annotated images, we evaluate two deep learning-based approaches for automated leaf counting: the first based on counting-by-regression from whole image analysis and a second based on counting-by-detection. Both approaches can achieve RMSE (root of mean square error) smaller than one leaf, only moderately inferior to the RMSE between human annotators of between 0.57 and 0.73 leaves. The counting-by-regression approach based on CNNs (convolutional neural networks) exhibited lower accuracy and increased bias for plants with extreme leaf numbers which are underrepresented in this dataset. The counting-by-detection approach based on Faster R-CNN object detection models achieve near human performance for plants where all leaf tips are visible. The annotated image data and model performance metrics generated as part of this study provide large scale resources for the comparison and improvement of algorithms for leaf counting from image data in grain crops.

## Introduction

Plant development, unlike that of many vertebrates, incorporates the potential for substantial plasticity in organ numbers. Different varieties of the same species may produce different numbers of leaves prior to flowering (Kim et al., 2017; Allen et al., 1973). Two genetically identical clones may produce different numbers of leaves prior to flowering when grown in different environments (Tollenaar and Hunter, 1983; Brooking et al., 1995). Variation in both the total number of leaves produced and the phyllochron, or the rate in thermal time at which leaves appear, are associated with a wide range of agricultural plant properties. For example, in maize, the number of leaves is correlated with plant height, flowering time, and moisture at harvest (Allen et al., 1973). In switchgrass, a larger number of leaves, indicative of a long duration of vegetative growth prior to reproductive transition, is highly correlated to the yield (Van Esbroeck et al., 1997). In potato, the number of green leaves has been used as indicator to determine drought resistant and susceptible varieties (Deblonde and Ledent, 2001). In perennial ryegrass, the number of leaves was used as a criterion for determining defoliation time (Fulkerson and Slack, 1994). Many crop growth models also incorporate both leaf appearance rate and final leaf number as crucial parameters which must be determined when these models are fit to new crops or new varieties of existing crops (He et al., 2012; Lizaso et al., 2003; Hammer et al., 2010; Truong et al., 2017). Whenever, even within individual genetically uniform crop varieties, canopy development and architecture of individual genetically identical crop varieties across locations and years in response to environmental factors. (Kumar et al., 2009) As a result, counting the number of leaves on individual plants is a task frequently undertaken by plant biologists, plant breeders, and agronomists.

However, manually counting leaves can be time consuming and logistically challenging. This is particularly true for large studies where hundreds of genotypes are being evaluated, each with many replicates (e.g., QTL mapping, GWAS, breeding trial evaluations, etc.). Leaves produced early in development can senesce and be undetectable by maturity (Figure 1A and B). As a result, multiple measurements throughout development are necessary not only to track leaf appearance rates but also to have an accurate final count of the total leaves produced. These challenges have limited the number of studies identifying genetic variants influencing the number of leaves or leaf appearance rates (Li et al., 2016; Cao et al., 2016; Hasan et al., 2016; Méndez-Vigo et al., 2010). The same challenges also hinder the development and deployment of genotype-specific crop growth models (He et al., 2012; Technow et al., 2015; Masjedi et al., 2018). These logistical hurdles have motivated the exploration of automated or semi-automated solutions to plant phenotyping (for example, (Furbank and Tester, 2011; Araus and Cairns, 2014; Fernandez et al., 2017; Zhang et al., 2017; Zhou et al., 2019). Advances in automated plant imaging have produced large plant image datasets (Liang et al., 2017; Feldman et al., 2017; Zhang et al., 2017). However, developing and deploying the most effective algorithms to allow the automated counting or scoring of the number of leaves from raw images of plants is an ongoing challenge within the plant phenotyping community.

**Figure 1.**
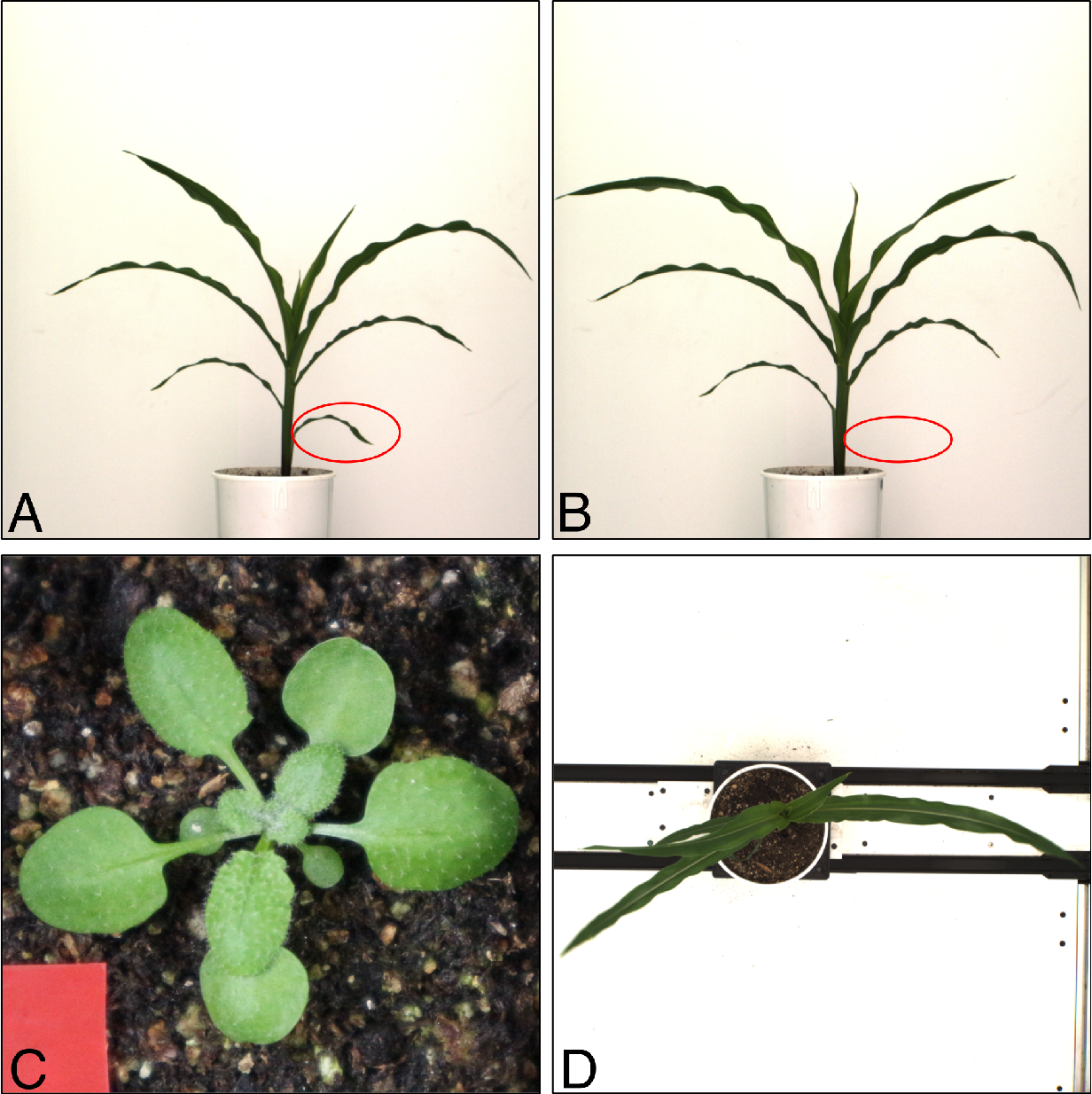
Challenging in tracking leaf number in plants. Leaves produced early in development can senesce, detach, and die over the lifetime of the plant, becoming undetectable at maturity. (A) A maize plant at 32 days after planting with eight visible leaves. (B) The same maize plant at 34 days after planting. The top three leaves have all lengthened significantly, while the bottom leaf – indicated in red ovals – has senesced and is no longer visible, resulting in only seven visible leaves for this older plant. (C) Illustration of the rosette leaf architecture exhibited by an A. *thaliana* plant in a top-down photo with all leaves visible. (D) Top-down photo of the same maize plant shown in panels A and B. Leaf occlusion in Panel D is more severe than in panel C.

A range of deep learning-based approaches to leaf counting have been proposed in recurring plant phenotyping workshops, conferences, and research articles (Aich and Stavness, 2017; Giuffrida et al., 2018a; Dobrescu et al., 2019). Many of these approaches are developed and validated using image and ground truth data from rosette plants (Giuffrida et al., 2016; Kuznichov et al., 2019). Rosette plants are defined by a shared leaf architecture where leaves are arranged in a compact circular pattern along the stem with substantially reduced stem elogation between leaves (Figure 1C). The focus on rosette plants is likely a result of the popularity of *A. thaliana*, a plant genetic model which belongs to the Brassicaceae, a family of plants including many species which grow in a rosette fashion. For plants exhibiting a rosette leaf architecture, it is frequently possible to collect images of the majority of non-occluded leaves from a single top-down view in a short period of time using relatively simple imaging systems (Scharr et al., 2014; Giuffrida et al., 2016, 2018a; Bell and Dee, 2019). Giuffrida and co-workers proposed and validated a unified deep network which could predict leaf numbers across multiple species which share a rosette-style leaf architecture (Giuffrida et al., 2018b). For the majority of species of monocots, leaves are sessile with the blade of the leaf directly joining the stem. Maize, wheat, and rice, – three monocot crops in the grass family – have elongated internodes between sessile leaves which emerge following alternating phyllotaxy. These crop species also provide roughly half of all calories consumed by humans around the globe (Tester and Langridge, 2010). Leaf counting has been shown to be practical in sorghum when using depth cameras to reconstruct 3D models of individual plants (McCormick et al., 2016; Xiang et al., 2019) However, only limited work has been conducted in leaf counting from 2D images of grain crops such as maize and sorghum (Pound et al., 2017; Zhou et al., 2020), potentially because of the lack of large, annotated, public training datasets for these crops. Unlike rosette plants, counting leaves from top-down images is not practical in these species (Figure 1D). Instead, the leaves of grasses are generally counted when viewed from the side which reduces the incidence of occlusion relative to top down views.

In this study, we assembled a maize image dataset using a non-invasive high throughput plant phenotyping platform (Ge et al., 2016). A set of 122,290 RGB images of maize plants, consisting of 10 images per plant per day from 923 unique plants representing 342 unique maize genotypes, were collected as part of two experiments conducted in 2018 and 2019. A parallel sorghum dataset of 27,770 images, consisting of five images per plant per day was collected from 343 unique plants representing 295 unique sorghum genotypes. The position and classification of individual leaf tips in a subset of these images were annotated by humans using the Zooniverse citizen science/crowd sourcing platform. Leaf counting approaches employing convolutional neural networks (CNNs) trained for regression and Faster R-CNN object detection models trained for leaf tip detection were evaluated. The release of raw data including leaf tip position and condition annotations, as well as the initial performance estimates of different models provide the community a benchmark to test against when seeking future improvements in automated leaf counting for grasses and grain crops.

## Results

### Assessment of the Leaf Annotation Dataset

Ten images for a maize plant were photographed from 10 different viewing angles each day (Figure S1A). Ground truth leaf numbers should not vary among images of the same plant taken on the same day, however apparent leaf numbers may vary as a result of differences in occlusion among images taken from different perspectives. The viewing angles of 108° and 288° were identified as the most straightforward to annotate, with leaves tending to have comparatively fewer occlusions or crossovers among leaves than other viewing angles. The images taken from the viewing angle of 108° were selected for the manual annotation. Annotators were asked to divide leaves into three categories: 1) leaves where the leaf tip was not visible (e.g. out of frame or occluded by another leaf); 2) leaves where the leaf tip was visible but mechanically damaged or cut; and 3) leaves where the leaf tip was visible and healthy (Figure S2). The total number of leaves for each annotated image was calculated by summing up all three leaf categories. The calculated number of leaves for the annotated image was also assigned to the other nine images with different viewing angles but captured for the same plant on the same day. Details of the annotation process and the definition of each of the three categories are provided in the methods section. Annotated leaf numbers for maize plants in this dataset ranged from 3 to 20 with reduced representation at both extremes (Figure S3A). All images and annotations are being released, but in the analyses below, we focus only on images of plants with between 4 and 15 leaves with each category containing at least 3,000 images of plants (Figure S3A).

A subset of 1,768 images were shown to multiple human observers. In 72% of cases both observers who saw a given image agreed on the total number of leaves (root mean square error (RMSE) = 0.71) (Figure 2A). Inconsistencies between observers were more common in images with larger total numbers of leaves. Manual follow-ups using the positional information recorded as part of the annotation process found that between observer disagreements were primarily explained by three issues: 1) differences when a new leaf, emerging from the top of the whorl, was included in the leaf count for the plant as shown in Figure 2B; 2) differences whether senescing or severely damaged leaves at the base of the stalk were still included in the leaf count for the plant (Figure 2C); and 3) partially occluded leaves which were missed by one observer but not another as shown in Figures 2D and E. Among assignments to the three different classes of leaves – healthy leaf tip, damaged leaf tip, and non-visible leaf tip – the one error we observed frequently was that at least some human annotators tended to annotate leaves which extended out of frame as healthy leaf tips rather than the non-visible leaf tips.

**Figure 2.**
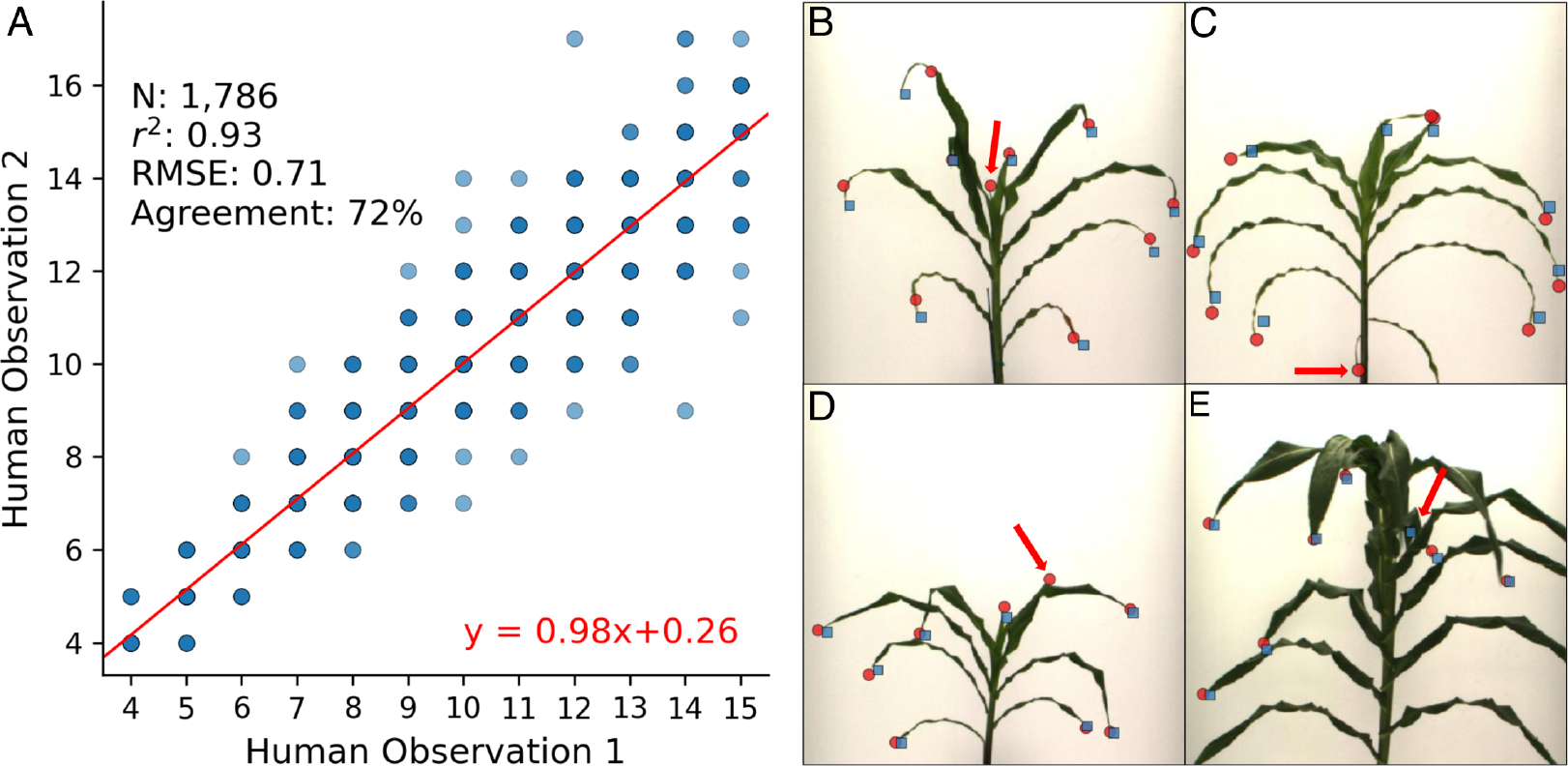
Human annotation errors on leaf counting. (A) Concordance in leaf number annotation between independent annotations of the same image by different observers. Dark blue points represent observations from larger numbers of distinct images while light blue points represent observations from smaller numbers of distinct images. The total tested images, r2 (square of the correlation coefficient), RMSE (root of mean square error), and the agreement rate are shown in the upper left corner. The best linear regression line is shown in red and the equation for that line is given in the bottom right. (B) An example of an image where two observers disagreed on the identification of a new leaf emerging from the whorl of a maize plant (red arrow). Selected positions of leaf tips for one observer are indicated with red circles and the other with blue squares. (C) An example of an image where two observers disagreed about the annotation of a senescing and potentially damaged leaf (red arrow). (D) An example of an image where one observer identified a partially occluded or overlapping leaf which was missed by a second observer (red arrow). (E) An example of a plant where phyllotaxy has shifted for upper leaves, making it quite difficult to annotate from any one side viewing angle. Images for panels B-E were cropped to aid viewing at a reduced figure size. Annotations which were made outside of the cropped portion of the image included in the figure are not shown.

Data leakage can occur when an annotated image dataset includes individual images sharing many common features and common labels, and these similar images are randomly partitioned into training and testing datasets. Data leakage can result in CNNs with overly optimistic prediction accuracy, i.e., the estimated results from CNNs are impossible to achieve in real world scenarios (Samala et al., 2020). At least two potential sources of data leakage exist in this leaf number/leaf position dataset. Firstly, this dataset includes images of the same plant collected only several days apart. These images will tend to look much more similar to each other than random pairs of images, and tend to contain plants with close or identical numbers of leaves. Secondly, variation in leaf number is under genetic control in maize (Cao et al., 2016), and this dataset includes images of multiple plants from the same genotype. Images of different, genetically identical individual plants will tend to be more similar to each other than images from random unrelated individual plants, and will be more likely to share close or identical numbers of leaves at the same stage in development. To avoid problems of data leakage, we elected to conduct data splitting at the genotype level, rather than splitting at the level of individual images or individual plants when preparing data for training and testing. All images from 50 genotypes were set aside as testing data to evaluate final model accuracy. The remaining 292 genotypes were split into five groups of training and validation datasets through five-fold cross-validation for both the counting-by-regression approach based on CNNs and the counting-by-detection approach based on Faster R-CNNs described below.

### Counting Leaves by Regression

Multiple deep learning based approaches have been applied to the task of counting the number of leaves in plant images (Pound et al., 2017; Dobrescu et al., 2019) Among them, the CNNs that treat the leaf counting task as a regression problem are most widely deployed for the direct estimation of leaf numbers. In this study, two different sets of training data were used when training CNNs to predict the correct number of leaves based on a single image of a maize plant. The first set of CNNs were trained using maize images with all 10 viewing angles in the training dataset. We acknowledge that even for human annotators, accurately counting leaves in images taken from some perspectives (e.g. viewing angles parallel to the plane of phyllotaxy) will be more challenging than others (e.g. viewing angles perpendicular to the plane of phyllotaxy) (Figure S1B). Therefore, a second set of CNNs were trained only using those images with the ‘best views’ in the training data. The best views were the two images where the distances between the left most and right most plant pixel were greatest among all 10 viewing angles, whereas the rest of eight images were treated as ‘other views’ (Figure S1B). The trained CNNs were then evaluated on the images, including all ten viewing angles from the 50 genotypes set aside as holdout test data. However, model performance was different when evaluated based on the testing images with different viewing angles.

For models trained using data from all 10 viewing angles, performance was significantly higher when evaluated using only the “best” viewing images than when evaluated using data from the “other” viewing images in the test data (Table 1, S1). Based on the greater prediction accuracy of CNNs when evaluating using only “best” viewing angle images, the decision was made to train a second set of CNNs employing only “best” viewing angle images as training data. The performance of these new models was comparable to the set of models trained using data from all 10 viewing angles when evaluating them on the “other” viewing images from the holdout test set. However, the new CNN models achieved a significantly greater correlation between prediction and ground truth (r^2^ = 0.87; p=0.011; t-test), higher agreement rate (45%; p=0.031; t-test), and lower RMSE (0.96 leaves; p=0.001; t-test) when tested on the “best” viewing images from the holdout test set (Table 1). The training time required for models trained using ‘best’ viewing images was also much shorter than the training time required for models trained using all 10 viewing images as the number of ‘best’ viewing images was only 1/5th of all viewing images.

**Table 1.** Performance of regression CNNs on leaf counting in maize

| Model | Testing data | $r^2$ | Agreement rate | RMSE |
| --- | --- | --- | --- | --- |
| CNN1 | Other Views | $0.79 \pm 0.04$ | $0.33 \pm 0.02$ | $1.28 \pm 0.11$ |
| CNN1 | Best Views | $0.84 \pm 0.03$ | $0.37 \pm 0.03$ | $1.11 \pm 0.09$ |
| CNN2 | Other Views | $0.78 \pm 0.02$ | $0.35 \pm 0.02$ | $1.31 \pm 0.12$ |
| CNN2 | Best Views | $0.87 \pm 0.01$ | $0.45 \pm 0.01$ | $0.96 \pm 0.04$ |

CNN1: CNNs trained using all 10 viewing angle images collected from each plant; CNN2: CNNs trained using only the two "best" viewing angle images.

Outcomes for the single highest performing model, one of the five trained using only the “best” viewing images, were r^2^ of 0.88, an agreement rate of 46%, and an RMSE of 0.92 (Figure 3A). For most numbers of ground truth leaves, the predictions of the CNN were not obviously biased towards predicting too many or too few leaves (Figure 3). However, the model did tend to overestimate the number of leaves for plant images with only four leaves, while tending to underestimate the number of leaves for plant images with 14 or 15 leaves (Figure 3A). These three ground truth values (4 leaves, 14 leaves, or 15 leaves) were the least common among leaf number groups included in our training dataset (Figure S3A). When excluding these three leaf number classes, the RMSE between model and annotator was 0.88 leaves, although this was still higher than the RMSE observed among human annotators (0.72 leaves) (Figure S4, 2A).

**Figure 3.**
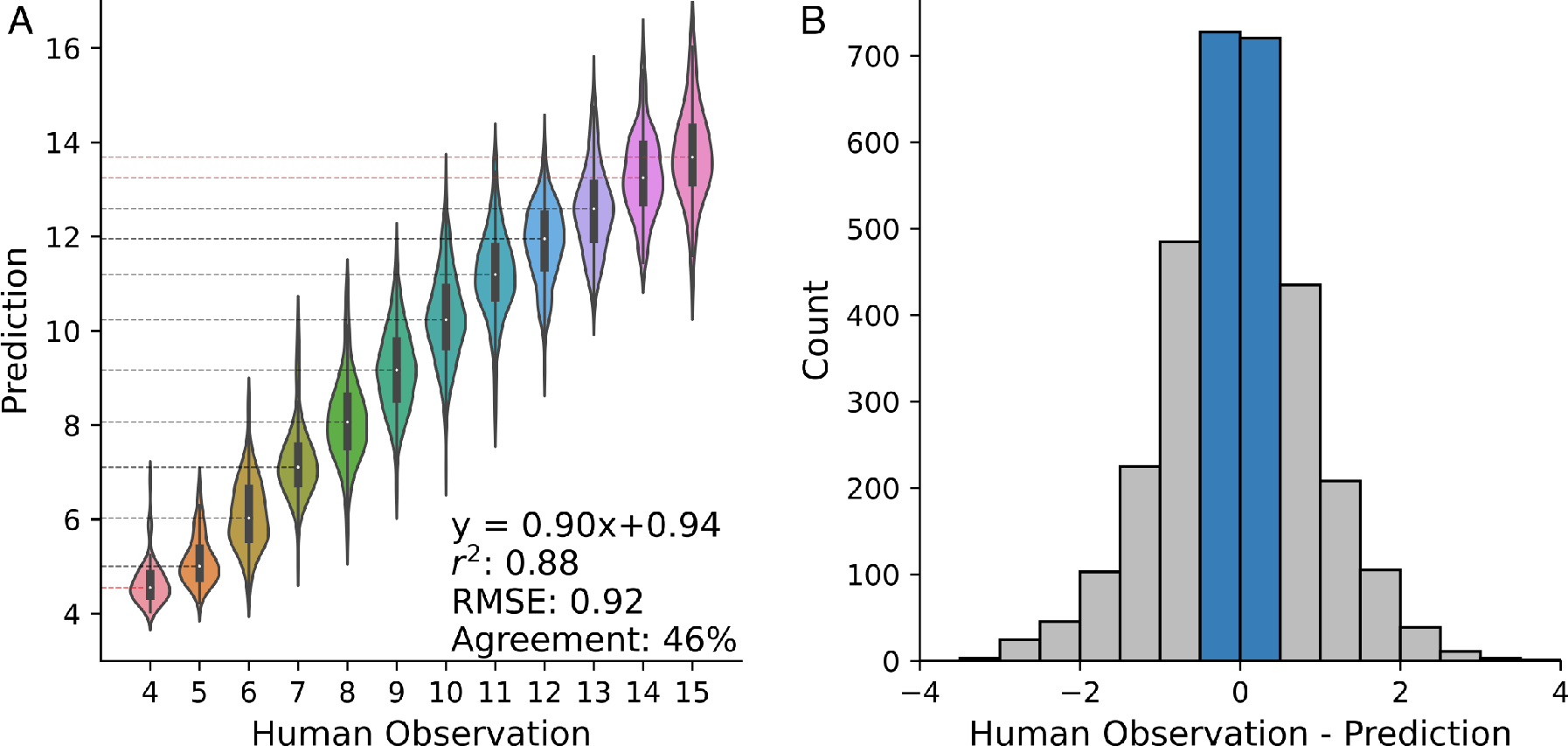
Distribution of predictions and errors for counting-by-regression approach. (A) Relationship between predicted leaf number and human annotated number of leaves among the “best” viewing images for the single best performing CNN trained using the “best” viewing images. The equation of the best linear regression, r2 (square of the correlation coefficient), RMSE (root of mean square error), and the agreement rate are indicated in the bottom right corner. Dashed lines indicate the median predicted leaf number values for images with each different number of leaves identified by human annotators. Cases where the median value for images of plants with a given number of leaves diverges from the annotated number of leaves by >0.5 leaves are indicated in red, all other cases indicated in black. (B) Distribution of error (difference between human observation and predicted number of leaves) for the same single best performing model shown in panel A. The absolute difference of less than 0.5 indicated by the blue bars was used for the calculation of agreement rate.

### Application of Maize Models to Sorghum

Maize and sorghum share similar plant architectures prior to flowering. Untrained observers can struggle to tell the difference between vegetative stage maize and sorghum plants, and even trained observers will struggle at the seedling stage. We generated annotations for a published set of sorghum images which were collected using the same automated imaging facility employed to generate the maize images described above (Miao et al., 2020). In the sorghum data, only five images were collected from five side viewing angles instead of ten in the maize data. The sorghum dataset annotated in this study exhibited a narrower distribution of leaf numbers than the annotated maize images (Figure S3B). At least 500 images were available of sorghum plants with between 6 and 13 leaves, and images of plants with these numbers of leaves were employed below. Models trained using ‘best’ viewing maize images and directly applied to sorghum exhibited a decline in performance relative to the performance of those same models in maize across all three metrics (Table 2). The r^2^ between annotated and predicted leaf numbers decreased to 0.54 from 0.87, the agreement rate decreased to 35% from 45%, and the RMSE increased to 1.14 from 0.96. The decline in r^2^ is likely explained in part by the reduced variability in leaf numbers among plants in the sorghum image dataset relative to the maize dataset (Figure S3B). However, this difference cannot explain the decline in the agreement rate or the increase in the RMSE. A second approach was evaluated where a subset of the annotated sorghum data was employed to retrain the last fully connected layer of the same adopted maize models. This transfer learning based approach resulted in small but significant improvements in all three evaluations of model performance. The mean RMSE of the new models through transfer learning was only 1/10th of a leaf less accurate in sorghum than the original models were in maize (Table 2).

**Table 2.**
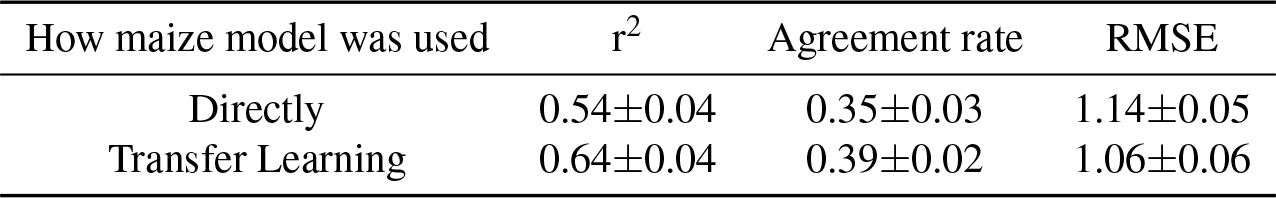
Performance of regression-CNNs on leaf counting in sorghum images

| How maize model was used | $r^2$ | Agreement rate | RMSE |
| --- | --- | --- | --- |
| Directly | 0.54±0.04 | 0.35±0.03 | 1.14±0.05 |
| Transfer Learning | 0.64±0.04 | 0.39±0.02 | 1.06±0.06 |

### Counting Leaves by Detection

Human beings possess and employ at least two mechanisms for counting objects: subitizing, the near instantaneous recognition of object numbers for sets of one through four, and a slower counting based mechanism for larger quantities (Trick and Pylyshyn, 1994). The regression-based approach described above is, in some ways, more analogous to subitizing as it seeks to determine the complete number of leaves present in the image at once. In any case, the counting-by-regression approach is sensitive to the number of images representing each potential number of leaves a plant can have, with biased performance for rarer and more extreme leaf numbers (Figure 3A, S4). An alternative approach to leaf counting is to implement counting-by-detection models (Xu et al., 2018; Buzzy et al., 2020) built on top of object detection frameworks (Redmon et al., 2016; Ren et al., 2015). Compared to a single number as the label for each training image in building CNNs, Object detection models are trained using annotations of bounding boxes around objects of interest.

In order to create bounding boxes centering on leaf tips as training data for leaf tip detection, the annotated coordinates of each leaf tip were expanded by 15 pixels in each direction resulting in a 30 x 30 pixel bounding box centered on each annotated leaf tip. The resulting bounding box annotations were employed to train a set of Faster R-CNN object detection models through five-fold cross-validation using the same splitting strategy employed above in maize. Each Faster R-CNN model employed the Resnet50 architecture as a backbone and was pre-trained on the ImageNet dataset (Russakovsky et al., 2015; Ren et al., 2015; He et al., 2016). All object detection models were trained on the bounding box annotations with intact and damaged leaf tips as two distinct objects. During the training process, model performance was evaluated using standardized metrics: average precision and recall for the purpose of stopping the training process early to avoid over fitting. Final trained models were evaluated using the same metrics used in evaluating CNNs by comparing the leaf numbers annotated by humans and predicted numbers of leaves by summing all detected leaf tips by object detection models. Tested was conducted using directly annotated images from plants in the 50 genotype hold out test set.

Unlike the CNN-based counting-by-regression approach where the number of leaves in an image from one view could be generalized to images taken at different viewing angles for the same plant on the same day, training object detection required information on the position of each leaf tip. As a result, training data consisting solely of directly annotated images collected from genotypes in the training set. Evaluation of performance was conducted using two subsets of the holdout test data. The first subset consisted of all directly annotated images collected from plants belonging to genotypes in the holdout test dataset. The second included only the subset of the images in set one where human annotators reported that all leaf tips were in frame and not occluded. When evaluated across all directly annotated images in the holdout test set, object detection models achieved an average r^2^ of 0.78, an agreement rate of 43% and an RMSE of 1.33 (Table 3). Manual examination of images where the predicted number of leaves was less than the annotated number of leaves revealed that the majority of missing detections resulted from leaves which were either out of frame or occluded by other leaves (Figure 4B). The latter problem – occlusion of leaves by other leaves – is likely unavoidable in 2D representations of 3D plants. However, leaf tips which are missed because the plant is wider than the imaging chamber could be addressed through changes to image acquisition (e.g. make a bigger imaging chamber for bigger plants like mature maize). Manual examination of images where the predicted number of leaves was greater than the annotated number of leaves indicated that the two most common sources of these errors were tassel branches being misclassified as leaves, and the tips of ear husk leaves being detected as leaves. Husk leaves are homologous to normal leaves in maize but traditionally not included when counting leaves in that species (Figure 4C).

**Table 3.** Faster-RCNNs Leaf Counting Performance

| Testing data | $r^2$ | Agreement rate | RMSE |
| --- | --- | --- | --- |
| Directly Annotated | $0.78 \pm 0.01$ | $0.43 \pm 0.01$ | $1.33 \pm 0.04$ |
| Directly Annotated - All Tips Visible | $0.88 \pm 0.01$ | $0.56 \pm 0.03$ | $1.00 \pm 0.06$ |

**Figure 4.**
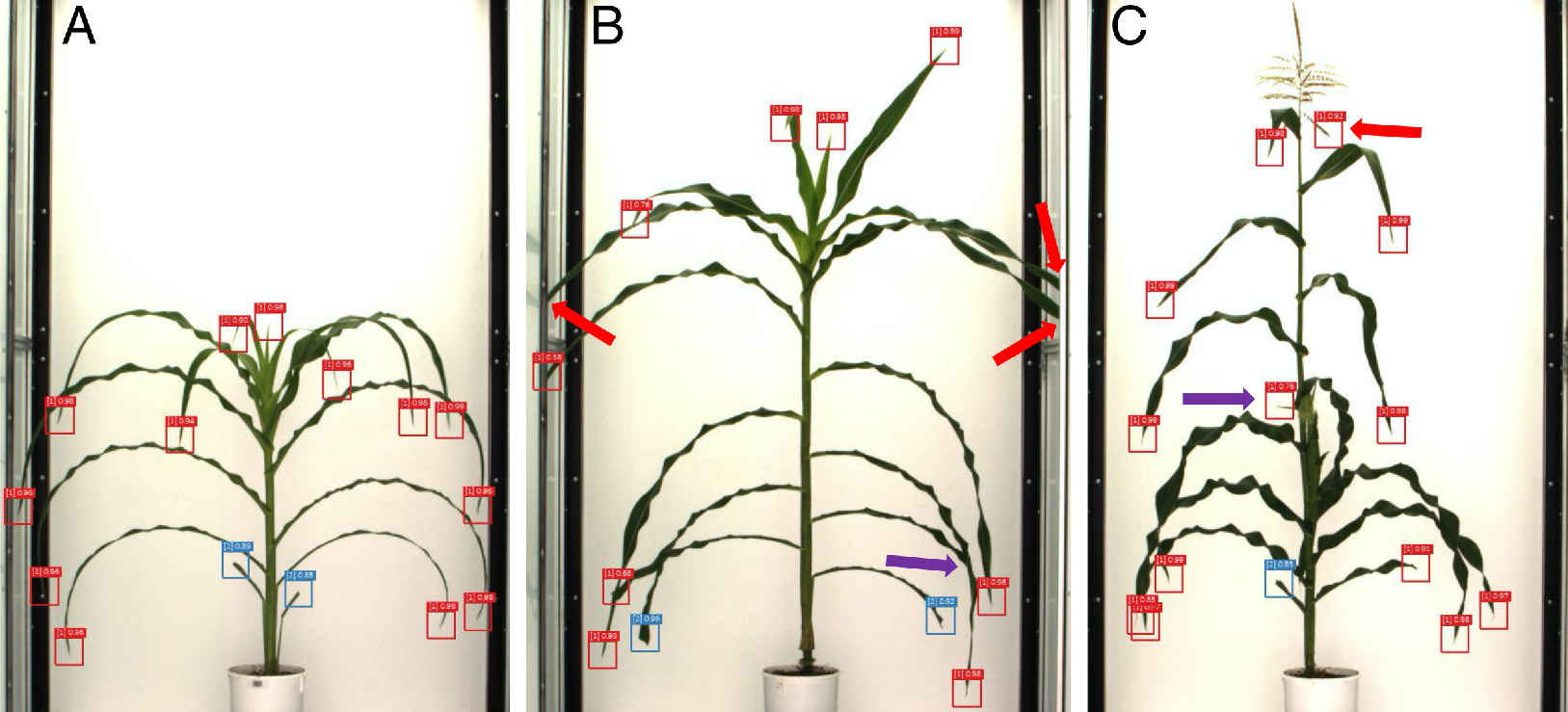
Representative leaf tip detection results. (A) Example of a plant with perfect detection, all annotated leaf tips are correctly identified. Undamaged leaf tips detected by the model indicated in red, damaged leaf tips detected by the model indicated in blue. The values above each bounding box indicate the classification confidence score assigned by the Faster R-CNN model. (B) An example of a plant where multiple leaves were missed by the object detection model either because the leaf tip itself is out of frame (red arrows) or because the leaf tip is occluded behind another leaf of the same plant from this viewing angle (purple arrow). (C) Example of a plant where two erroneous leaf tips were detected by the object detection model. One erroneous detection is centered on a tassel branch (red arrow), although the majority of tassel branches are not misclassified as leaves. The second erroneous detection is centered on a husk leaf associated with a developing ear on this particular maize plant (purple arrow).

When trained object detection models were evaluated solely on the subset of images where all leaf tips were reported to be visible by human annotators, the performance of these models increases substantially with a mean r^2^, agreement rate, and RMSE of 0.88, 56%, and 1.0 respectively (Table 3). The best Faster R-CNN as part of the five-fold cross-validation process had an r^2^ of 0.90, and agreement rate of 59.5% and an RMSE of 0.92, which is competitive compared to human performances on the same type of testing images in maize (Figure 5).

**Figure 5.**
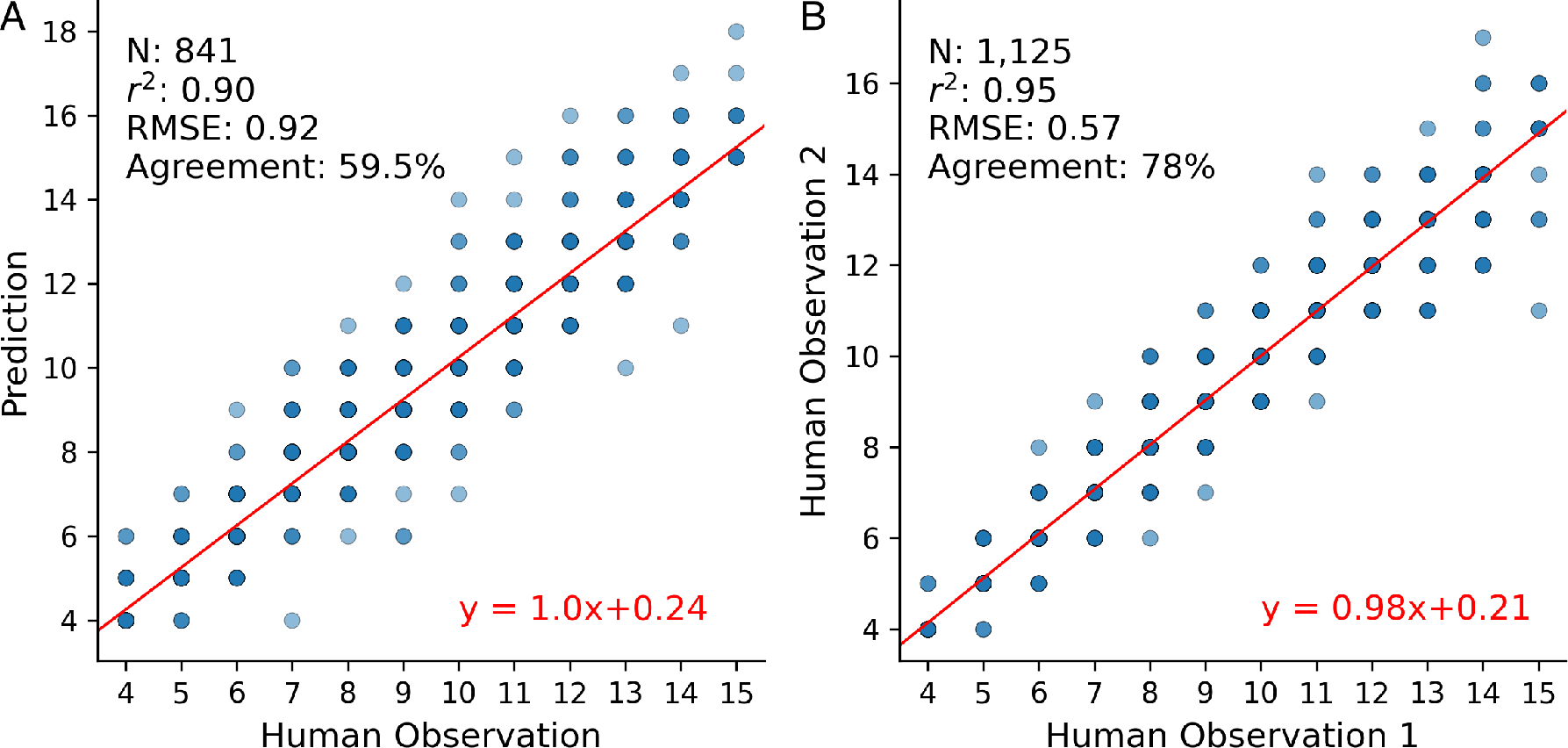
The performance of the best object detection model on images without leaf tip occlusions. The performance of the best object detection model on images without leaf tip occlusions. (A) Relationship between predicted leaf number and human annotated leaf number among directly annotated images without leaf tip occlusions for the single best performing object detection model. (B) The concordance in leaf number annotation between independent annotations on the images without leaf tip occlusions. The number of tested images, r^2^ (square of the correlation coefficient), RMSE (root of mean square error), and the agreement rate are indicated in the upper left corner. The best linear regression line and the corresponding equation were indicated in red color.

## Discussion

In this study we generated an annotation benchmark dataset for leaf counting and detection in two grain crops: maize and sorghum. The annotations capture both the number of leaves and the position of the leaves in each image. In total, 118,445 leaves were directly annotated in 12,229 maize images, and 53,247 leaves were directly annotated in 5,554 sorghum images. Because our dataset includes images of the same plant taken from different viewing angles and the true number of leaves a plant possesses is fixed at any given point at time, we are able to generalize information on leaf numbers, but not position, to other images of the same plants taken on the same day from different viewing angles. The results are a total of 122,290 images of maize plants and 27,770 images of sorghum plants with ground truth leaf number annotations. These annotations are not perfect. Comparison of annotations of the same images by multiple human observers found that observers agree on the number of leaves for 72% of images, with an overall RMSE of 0.71 leaves. To establish initial benchmarks for performance on leaf counting in this dataset, we evaluated two approaches based on deep learning models: a CNN-based counting-by-regression approach and a counting-by-detection approach using Faster R-CNN object detection models. Both approaches were able to achieve RMSE values of less than one leaf, but exhibit different tradeoffs.

For the counting-by-regression approach, a CNN trained to output a floating point prediction of the number of leaves in a given image performed better on images where the plane of phyllotaxy was perpendicular to the viewing angle of the camera (Figure S1; Table 1). Similar challenges were found among human annotations when the plane of phyllotaxy was parallel to the viewing angle of the camera (Figure Figure 2E). We found a crude metric, the distance from the left most to the right most plant pixel in an image, was an effective approach to identify these images that are most amenable to leaf counting. CNN models trained with all images or only these two “best” viewing images per plant both exhibit better performance on these “best” viewing images, matching the performance they exhibited when assessed using only the subset of images which were directly annotated (Table 1, S1). There are two straightforward approaches which could be taken to further improve model performance: new and more optimized model architectures, and further improvements in the size and/or accuracy of labelled training data. The first depends on more advanced models proposed by machine learning experts. Records for leaf counting accuracy of rosette plants continue to be broken as new and better performing models and approaches are proposed and tested (Aich and Stavness, 2017; Ubbens et al., 2018; Dobrescu et al., 2020). The second depends on the availability and training of human annotators. We identified a number of common inconsistencies between different annotators (Figure 2B and C) which could be incorporated into training for subsequent rounds of annotation of images from maize, sorghum and other grain crop species. However, the counting-by-regression approach to predicting the number of leaves from CNNs faces another challenge which is somewhat harder to address. For many quantitative genetic and breeding applications the extremes of any phenotypic distribution will be of greatest interest, yet a key drawback of the CNN-based counting-by-regression approach is that performance declined for plants with the smallest or largest numbers of leaves in our dataset (Figure 3A, S4). From a computational perspective, this issue can be easily addressed by collecting more images of plants with these numbers of leaves and including them in the dataset. Biologically this approach presents additional challenges. Images of plants with fewer leaves can be collected by starting imaging earlier in development. However, after accounting for the senescence of juvenile leaves, only a small subset of maize genotypes in our study will carry 14+ leaves at any given time point. Additional replication of these genotypes could be included but this would create a bias in the training data with high leaf number plants belonging to a small number of genotypes which exhibit additional genetically controlled differences in appearance. Alternatively, data could be generated using photoperiod sensitive tropical maize lines grown under long day conditions which prolong vegetative development, resulting in the production of greater numbers of leaves (Stevenson and Goodman, 1972). Yet this approach would again mean that genetically and phenotypically distinct lines would represent different portions of the potential range of the total number of leaves observable in maize.

In contrast to the CNN-based counting-by-regression approach, the performance of the counting-by-detection approach using leaf tip detection models was consistent across plants with different numbers of leaves. The performance of this approach was modestly poorer than the CNN predictions across all directly annotated images, but achieved higher agreement rates with human annotations and comparable r^2^ and RMSE for images without hidden leaf tips (Table 1; Table 3). For images without any hidden leaf tips, the divergence between the annotation of different humans looking at the same images as quantified by RMSE was only 1/3 of a leaf better than the divergence between the best object detection-based leaf counting model and human annotation (Figure 5). False negative leaf detections primarily result from non-visible leaf tips, while false positive detections primarily result from husk leaves or tassel branches. Including ear shoot/ear and tassel as additional features in subsequent annotations of this or other image datasets would allow these organs to be included as additional objects when training object detection models. This would presumably reduce false positive detections. Leaf occlusions as a result of maize plants which were too large to fit entirely within the field of view of the imaging chamber are a specific constraint of the facility employed in this study (Ge et al., 2016), and are best addressed at the data acquisition level rather than data analysis. However, leaf overlap and self-occlusion will be an inherent problem whenever a 3D plant is represented as a 2D image. A similar object detection model was recently employed to estimate leaf number by detecting whole leaves in maize images (Zhou et al., 2020). However, this approach also suffers from issues with occlusion. Performance was reported using mean ± standard deviation of absolute difference between the predicted and ground truth leaf numbers instead of RMSE. For comparison purposes we calculated the same metric for our best performing leaf tip detection model. The best leaf tip detection model from this study achieved a value of (1.31±1.60) compared to value of (1.60±1.63) reported by Zhou and coworkers. However, this comparison should be treated with caution given the different images used for model evaluations and differences in training data size. By making all of our raw images and annotations publicly available (see Data Availability Statement) we hope to reduce the barriers to head-to-head comparison of different approaches to leaf counting performance in maize, and thus stimulate future improvements in performance. The frequency of self-occlusions is likely to vary among genotypes as both leaf curvature and phyllotaxy are under partial genetic control in maize (Ford et al., 2008; Giulini et al., 2004). Leaf occlusion is therefore likely to introduce heritable genetic error (Liang et al., 2017) if counting-by-detection is employed to phenotype maize plants in quantitative genetic or plant breeding projects. Transitions to methods which incorporate 3D information (McCormick et al., 2016; Xiang et al., 2019; Gaillard et al., 2020) and/or the development of approaches to integrate information on leaf tips detected in multiple images of the same plant taken from different angles will likely be ultimately needed.

The availability of leaf counting models which can match or exceed human performance would provide significant benefits to plant breeders, plant geneticists, and crop modelers. The number of leaves is employed as a morphological indicator in predicting flowering time and yield (Allen et al., 1973; Van Esbroeck et al., 1997). Many plant geneticists seek to identify the genomic regions and genes controlling natural variation in leaf number among members of the same species. For example, a study was able to map QTL controlling variation in the number of leaves in maize from manually scoring 866 inbred lines from a maize-teosinte BC2S3 population with five plants of each inbred scored in each of two years, requiring counting the leaves of at least 8,660 individual maize plants (Li et al., 2016). Automated leaf scoring would enable similar studies to incorporate more genotypes, include additional replication, or conduct a study of equivalent scale using fewer resources. Actual automated scoring of leaf numbers, combined with existing high throughput phenotyping facilities which image plants throughout development, would make it feasible to collect time series data for determining total leaf number and tracking leaf appearance rate, and monitoring how these values vary across different genotypes or in the same genotype grown under different environmental conditions. Studies of environmental stresses and genotype by environment interactions in crop species can translate to agricultural applications most directly when the studies themselves are conducted in the field. Yet a substantial gap in image analysis remains between the approach described in this study for analyzing data from individual plants photographed against an uncluttered background in a chamber with constant illumination, and models which could conduct accurate leaf counting under field conditions with varying illumination. Here, rosette plants again possess a significant structural advantage as field data is easier to acquire when imaging from the top down relative to the side. Top-down imaging tends to place the plant against a non-uniform but contrasting background of soil (Cao et al., 2016). Automated approaches for collecting side view images in the field are improving (Fernandez et al., 2017; Young et al., 2019); however side views tend to result in plants being photographed against a non-contrasting background of other plants in the same field.

## Methods

### Image Acquisition and Annotation

Maize image data was taken from two separate experiments conducted at the University of Nebraska-Lincoln’s Greenhouse Innovation Center (Latitude: 40.83, Longitude: −96.69) in 2018 and 2019. In the first experiment, 256 lines from the Buckler-Goodman inbred association panel (Flint-Garcia et al., 2005) were planted on July 23rd, 2018. In the second experiment, 255 lines from the Buckler-Goodman inbred association panel and 33 enhanced lines from the GEM (Germplasm Enhancement of Maize) project (Pollak, 2003) were planted on March 25th, 2019. Maize kernels were sown in 2.4 gallon pots with Fafard germination mix supplemented with 1 cup (236 mL) of Osmocote plus and 1 tablespoon (15 mL) of Micromax Micronutrients per 2.8 cubic feet (80 L) of soil. The target photoperiod was 14:10 with supplementary light provided by light-emitting diode (LED) growth lamps from 07:00 to 21:00 each day. The target temperature of the growth facility was between 24-26°C. After growing in the greenhouse for 28 days in the first experiment and 38 days in the second experiment, all the plants were moved to a high throughput phenotyping facility equipped with a plant conveying system and different kinds of imaging chambers (Ge et al., 2016; Gaillard et al., 2020). The conveyor belt transferred each pot to a set of imaging chambers once every two or three days for imaging, and to a watering station each day. At the watering station plants were weighed and watered back to a target weight to ensure all plants were growing in a good condition. In the imaging chamber, each plant was rotated 36 degrees nine times and an image was captured for each stop. Therefore, a total 10 photos with 0°, 36°, 72°, 108°, 144°, 180°, 216°, 252°, 288°, and 324° viewing angles were captured for each plant on the same day (Figure S1A). Plants were photographed from 28 to 73 DAP (days after planting) in the 2018 experiment, and 38 to 70 DAP in the 2019 experiment. These imaging time periods covered both late vegetative growth and flowering stages for the majority of genotypes in the population. A total of 122,290 maize images were acquired from the two experiments and each image had a resolution of 2,454 x 2,056 pixels.

Sorghum image data was taken from a previously published image dataset deposited in CyVerse (https://doi.org/10.25739/p39bdz61) by Miao et al. (Miao et al., 2020). This dataset consisted of 27,770 images collected from 343 unique sorghum plants representing 295 inbred lines from the sorghum association panel (Casa et al., 2008). Plants were photographed from July 26 to August 31, 2017 over a period of 37 days spanning vegetative and reproductive development for the majority of genotypes in the population. In contrast to 10 viewing angles in the maize dataset, on each imaging date sorghum plants were photographed from only 5 different viewing angles including 0°, 36°, 72°, 108°, and 144°.

As images with different viewing angles for the same plant captured on the same day share the same leaf number, only images taken from the 108° viewing angle were uploaded to the crowd sourcing platform Zooniverse for annotation. All the leaves in each image were grouped into three classes: 1) leaves with visible healthy leaf tip; 2) leaves with visible but damaged or cut leaf tip; 3) leaves with leaf tips that are not visible due to occlusions. For the first two classes where leaves had visible leaf tips, annotators were asked to click corresponding leaf tip positions. For leaves whose leaf tips were not visible, annotators were asked to click anywhere in the leaf (Figure S2). The total number of leaves in each image was calculated as the sum of the number of annotations for each of these three classes. We estimate it took approximately 132 total man hours to complete all annotations for 12,229 maize images and 5,554 sorghum images. All the annotation results including the coordinates of tip positions and the leaf numbers have been deposited in FigShare (See Data and Code Availability Statement).

### Image Processing

All images were preprocessed by cropping to a maximum bounding rectangle containing all plant pixels as shown in Figure S1. Removing uninformative parts in the image such as imaging chamber frames and the bottom pot made the image contain a higher proportion of plant pixels, which made it easier for deep learning models to learn plant related features. A pixel was classified as a plant pixel if the pixel value in the green channel was larger than 130. This cutoff was set after manually checking the performances of various threshold values on images across different genotypes and time points. Among the 10 images collected from a single maize plant on the same day, the two images where the distance between the furthest left and furthest right plant pixels was widest were included in the “best views” category and the other eight images were treated as ‘other views’ for downstream analyses (Figure S1B).

The preprocessed images were split to training, validation, and testing images at the genotype level, rather than at the plant or image level. This practice was adopted to avoid issues of data leakage which can occur when very similar images appear in both training and testing processes (Samala et al., 2020). All images from a set of 50 maize inbred lines generated as part of this study were set aside as testing data for evaluating the final accuracy of models. Image data for the remaining 292 lines were split for five-fold cross-validation with five sets of training and validation data. The first two sets contain all images from 233 inbred lines in the training dataset and all images from 59 maize inbred lines in the validation dataset. The remaining three sets contain all images from 234 and 58 inbred lines in training and validation datasets respectively.

### Training and Evaluation of CNNs

Two sets of maize CNNs were trained in this study: CNNs trained on all 10 viewing images, and CNNs trained only on the best viewing images (Figure S1). In each case, five models were trained through five-fold cross-validation. All the CNNs were trained using transfer learning by fine-tuning a pre-trained Resnet18 model (He et al., 2016) to fit customized maize images for leaf counting tasks. The pre-trained Resnet18 model adopted in this study was trained on the 1,000-class ImageNet dataset (Russakovsky et al., 2015) with 1,000 output features in the last fully connected layer. In order to adapt the model to the task of leaf counting, the number of output features was set to 1 representing one estimated leaf number for each input image. All the preprocessed images were resized to 224 x 224 pixels as the input image size during training and evaluating CNNs. In addition, two types of image transformation approaches were applied on the input training images. The first transformation approach changes input images by horizontally flipping images randomly with a given probability of 0.5. The second transformation approach randomly changes the brightness, contrast and saturation of the input images. The MSE (Mean Square Error) was used as the loss function considering leaf counting as a regression problem. The SGD optimizer was used with the ‘learning rate’ set to 0.001 and ‘momentum’ set to 0.9. The maximum epoch was set to 500 and early stopping was applied during the training with the ‘patience’ argument set to 50. This setting means that training will stop early if the overall MSE fails to improve for 50 continuous epochs. In this case the model with the smallest loss value will be saved and used for downstream evaluations.

A set of sorghum CNN models were also trained through transfer learning based on the CNNs trained on the best view maize image dataset. However, a different transfer learning strategy was adopted. Instead of fine tuning all the pre-trained weights, only the weights in the last fully connected layer were retrained using the best view sorghum images. All the sorghum images went through the same prepossessing steps as the maize images, including the removal of uninformative portions of each image. Sorghum images were also split on the genotype level to avoid data leakage, with all best view images from 250 sorghum lines used for training and all best view images for the remaining 44 lines used for testing. The hyperparameters for retraining sorghum models were the same as in training maize models.

Three metrics were adopted in this study to evaluate model performances: the square of the correlation coefficient (r^2^), the root mean square error (RMSE), and the agreement rate defined by the proportion of perfect predictions. A perfect prediction was called when the difference between ground truth and prediction was less than 0.5 so rounding would produce the same predicted integer value for leaves as reported ground truth. The mean and standard deviation of each metric were calculated from the five-fold cross-validation for the eventual model comparisons. The code for training and evaluating CNNs has been deposited on GitHub (See Data and Code Availability Statement).

### Training and Evaluation of Faster R-CNNs

A pre-trained Faster R-CNN object detection model (Ren et al., 2015) with Resnet50 architecture as the backbone was adopted to detect both healthy and damaged leaf tips in maize RGB images. The backbone was pre-trained on the ImageNet dataset and used as fixed feature extractor in Faster R-CNN. The Faster R-CNN model was pre-trained on the COCO 2017 instance dataset which includes 91 “stuff categories” including person, vehicles, fruits, and animals. To fit our leaf tip dataset, the output number of classes was set to three representing healthy leaf tip, damaged leaf tip, and background classes. Five object detection models were trained through five-fold cross-validation with the same splitting strategy used for CNNs, but only the directly annotated images were considered because the leaf tip positions were only annotated in these images. Each leaf tip coordinate indicated by (x, y) in a directly annotated image were converted to a bounding box coordinate (x-15, y-15, x+15, y+15) with a 30×30 pixel square centered around the leaf tip. The images used for both training and testing were the original plant photos without cropping as shown in Figure S1A. The SGD optimizer was used during training with the ‘learning rate’ set to 0.001, ‘momentum’ set to 0.9 and ‘weight decay’ set to 0.0005. The maximum epoch was set to 200 and early stopping was applied when both the average precision (AP) and average recall (AR) were converged. The object detection model was evaluated in the directly annotated images from the 50 genotypes set aside as holdout test data. The number of leaves estimated by the model was the sum of all the detected leaf tips with a classification score higher than 0.5. The code for training and evaluating Faster R-CNN models has been deposited on GitHub (See Data and Code Availability Statement)

## Acknowledgments

This work was supported by a University of Nebraska Agricultural Research Division seed grant to JCS, a National Science Foundation Award (OIA-1557417) to JCS, JY, and YG, and by the U.S. Department of Energy, Office of Science, Office of Biological and Environmental Research Program under award number DE-SC0020355 to JCS and AMT. This project was completed utilizing the Holland Computing Center of the University of Nebraska, which receives support from the Nebraska Research Initiative. This publication uses data generated via the Zooniverse.org platform, development of which is funded by generous support, including a Global Impact Award from Google, and by a grant from the Alfred P. Sloan Foundation. We thank Leighton Wheeler, Nate Pester, Isaac Stevens, Elijah Frost, Thomas Hoban, Sierra Conway, Elliot Kadrofske, Emma Chrzanowski, Carolina Freitas, Lou Townsend, and Logan Duryee for the work of image annotations in summer 2020. We thank Rebecca Roston and Jeremy Brown for gifting the Arabidopsis photo used in Figure 1.

## Data and Code Availability

All the maize and sorghum images as well as the corresponding leaf number and leaf tip position annotations used in this study have been deposited in Figshare: 1) maize leaf tip pixel annotations and images (https://doi.org/10.6084/m9.figshare.13056524.v1); 2) maize leaf number images and annotations (https://doi.org/10.6084/m9.figshare.13056512.v1); 3) sorghum leaf number images and annotations (https://doi.org/10.6084/m9.figshare.13056548.v1); 4) sorghum leaf tip images and annotations. (https://doi.org/10.6084/m9.figshare.13056527.v1). The scripts and source code related to training and evaluating CNN and Faster R-CNN models have been deposited on GitHub (https://github.com/freemao/MaizeLeafCounting).

## Supplementary materials

**Table S1.** CNN2 leaf counting performance on images collected from different viewing angles

| Viewing Angle | Proportion Classified as "Best" | $r^2$ | Agreement Rate | RMSE |
| --- | --- | --- | --- | --- |
| 288° | 0.27 | $0.88 \pm 0.01$ | $0.44 \pm 0.01$ | $0.95 \pm 0.04$ |
| 108° | 0.19 | $0.88 \pm 0.01$ | $0.44 \pm 0.01$ | $0.95 \pm 0.04$ |
| 252° | 0.14 | $0.84 \pm 0.01$ | $0.40 \pm 0.01$ | $1.12 \pm 0.07$ |
| 324° | 0.12 | $0.84 \pm 0.02$ | $0.38 \pm 0.02$ | $1.12 \pm 0.09$ |
| 72° | 0.09 | $0.84 \pm 0.02$ | $0.40 \pm 0.02$ | $1.11 \pm 0.09$ |
| 144° | 0.08 | $0.83 \pm 0.02$ | $0.38 \pm 0.02$ | $1.12 \pm 0.08$ |
| 180° | 0.03 | $0.71 \pm 0.04$ | $0.30 \pm 0.02$ | $1.57 \pm 0.17$ |
| 216° | 0.03 | $0.76 \pm 0.04$ | $0.32 \pm 0.03$ | $1.40 \pm 0.16$ |
| 0° | 0.02 | $0.70 \pm 0.04$ | $0.30 \pm 0.02$ | $1.55 \pm 0.17$ |
| 36° | 0.02 | $0.76 \pm 0.02$ | $0.33 \pm 0.02$ | $1.40 \pm 0.14$ |

**Figure S1.**
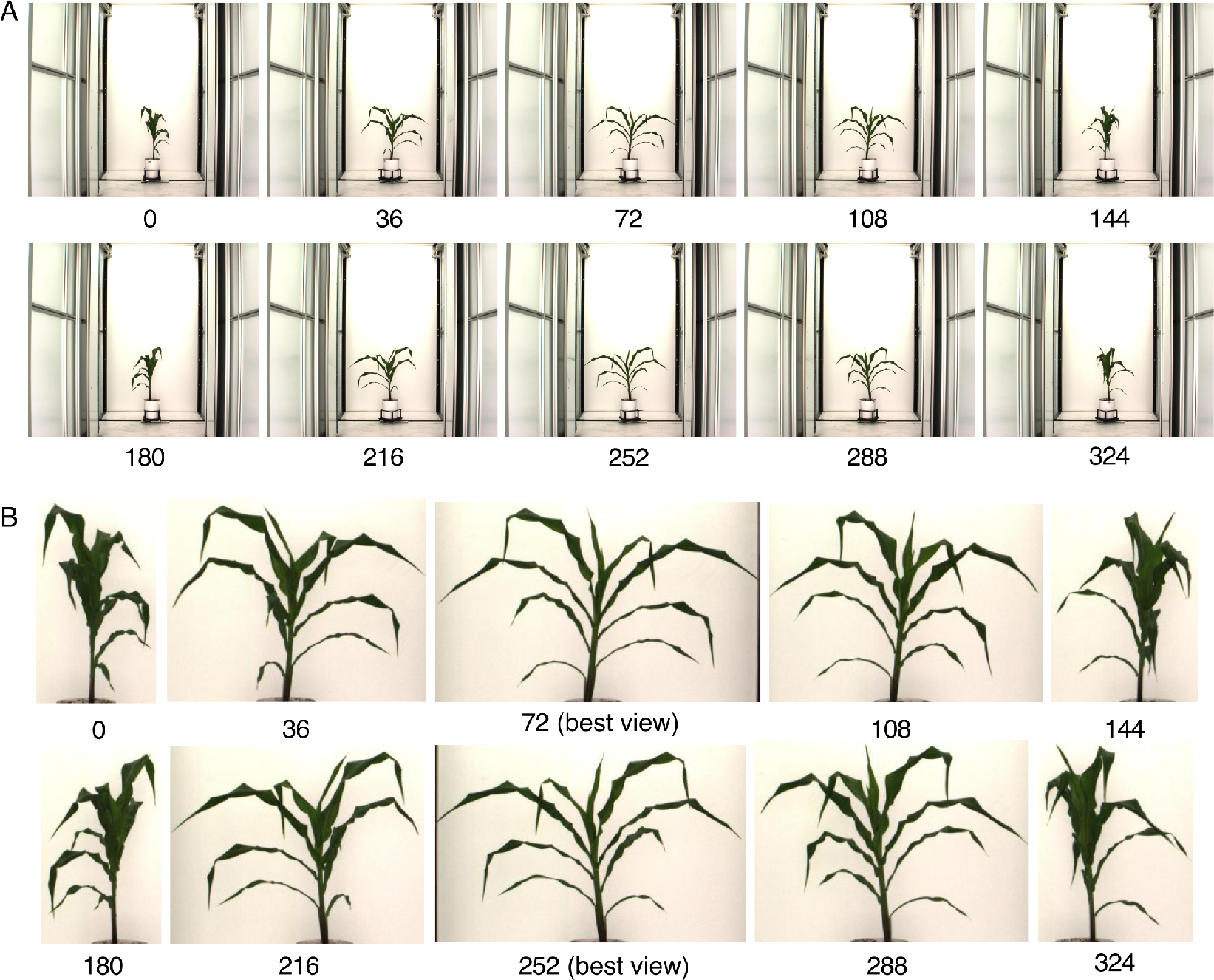
Characteristics of the images employed for annotation, training, and testing. Each maize plant was imaged sequentially 10 times, with 36 degrees of rotation between each image. Panel A shows the raw images generated by the automated phenotyping facility. Leaf counting was conducted using complete, uncropped images, with users having the capacity to zoom in on any given image to resolve finer details. Panel B shows the same images after each image is zoomed and cropped to a bounding rectangle containing all plant pixels. The two images with the widest bounding rectangles were classified as the “best” viewing angles and tended to be offset by five images, 180 degrees of rotation. Images in panel B were the images employed as training, validation, and testing data.

**Figure S2.**
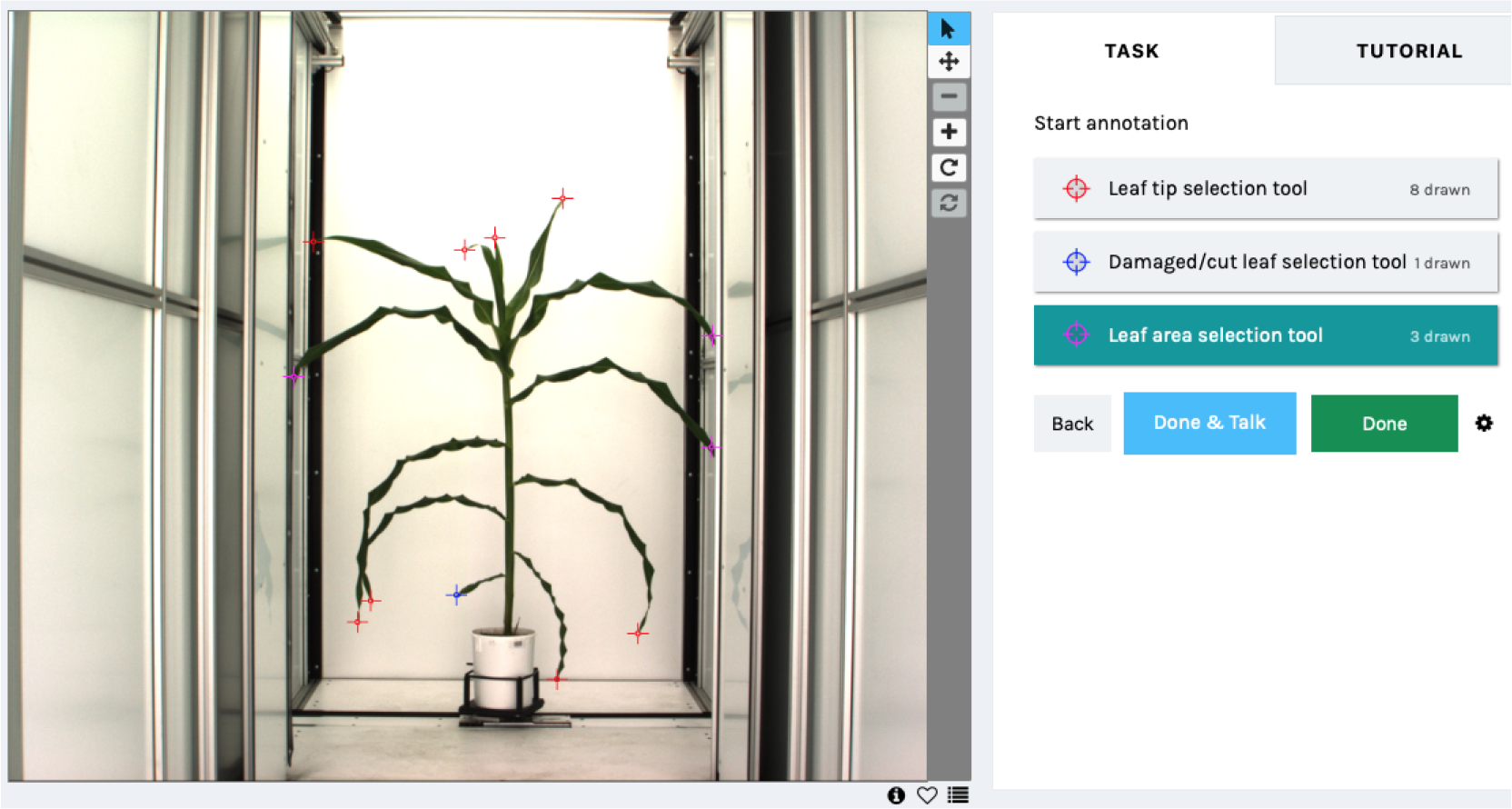
Example of the graphical interface on Zooniverse employed for leaf annotation. Users placed three different colors of markers - red, blue, and purple - to identify healthy leaf tips, damaged leaf tips, and leaves which were visible but where the leaf tip was out of frame or occluded, respectively. The total number of leaves present in a given image was the sum of the number of these three classes of annotation.

**Figure S3.**
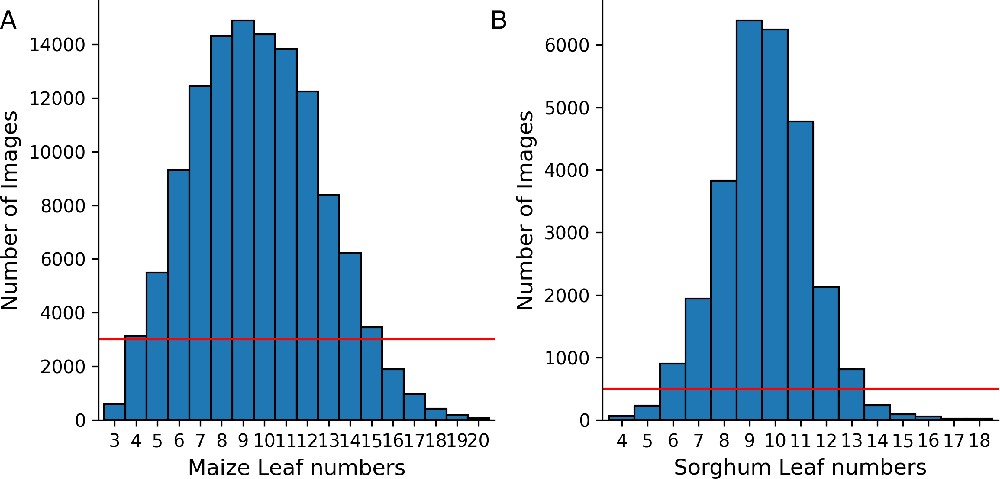
The distribution of observed leaf numbers among annotated maize and sorghum images. (A) Distribution of observed leaf number among maize images. Each annotated maize image informed the ground truth leaf number for a total of ten images collected on the same plant on the same day from different viewing angles. The red horizontal line indicates the cutoff of 3,000 images which was employed to select the range of leaf numbers employed in this study. (B) Distribution of observed leaf number among sorghum images. Each annotated sorghum image informed the ground truth leaf number for a total of five images collected on the same plant on the same day from different viewing angles. The red horizontal line indicates the cutoff of 500 images which was employed to select the range of leaf numbers employed when evaluating the performance of maize models on sorghum data.

**Figure S4.**
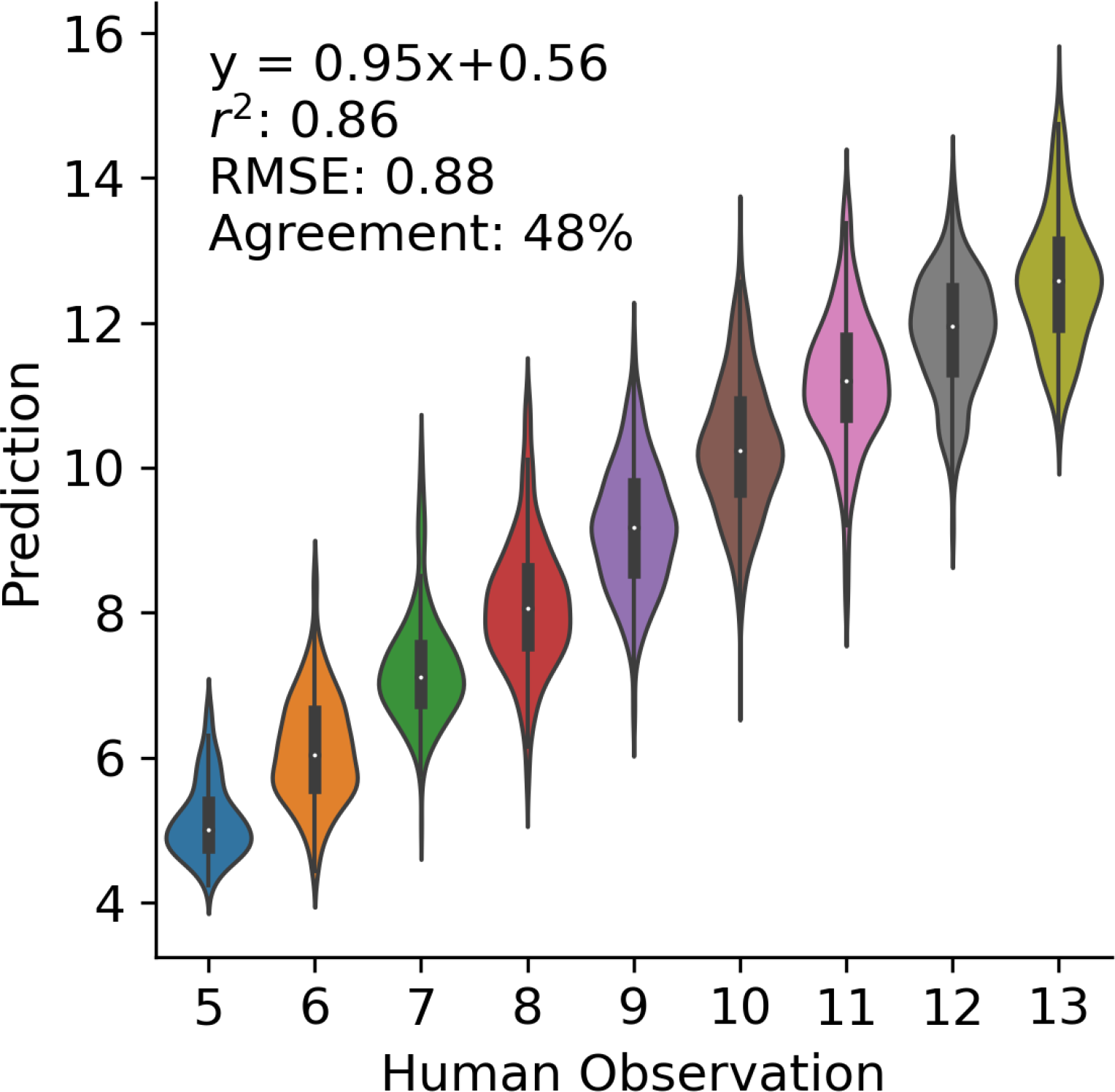
The performance of the best performing regression CNN when considering only images of plants with between 5 to 13 annotated leaves. The equation of the best liner regression, square of the correlation coefficient (r^2^), root mean square error (RMSE), and the agreement rate are indicated in the upper left corner.

